# Bayesian Stacking of Quantitative Adverse Outcome Pathways Isolates Primary Drivers of Liver Steatosis

**DOI:** 10.64898/2026.10.20.760409

**Authors:** Anna Melina Steinbach, Philip Marx-Stölting, Christian Tobias Willenbockel

## Abstract

The quantification of adverse outcome pathways (qAOPs) involving multiple converging pathways remains a fundamental challenge in toxicology. This difficulty is worsened by small in vitro datasets with missing values and heteroscedasticity, which limit the estimation of non-linear response functions and the robust estimation of uncertainty. To overcome these limitations, we developed a framework for estimating a Bayesian qAOP Ensemble (Bayesian qAOPE) that combines heteroscedastic dose-response functions with continuous splines and linear models. By combining multiple converging non-linear pathways through Bayesian stacking, the most meaningful pathways are identified, and strongly correlated responses are naturally accounted for. Applying our approach to a hepatic steatosis dataset, we demonstrate by exact Bayesian leave-one-out cross-validation that our framework yields an exceptional predictive performance suitable for the derivation of a probabilistically weighted Benchmark Dose (BMD). Our framework is highly adaptable and provides a chemical-agnostic, mathematically founded blueprint for integrating multiple converging pathways in a Bayesian network.

## 1. Introduction

The “Adverse Outcome Pathway” (AOP) approach has significantly expanded the field of toxicology by linking risk assessment based on in vivo adverse effects to mechanistic, biological sequences within the cell (Ankley et al., 2010; Villeneuve et al., 2014; Vinken, 2013). However, the transition from qualitative AOP networks to quantitative AOP networks (qAOPs) (Conolly et al., 2017) remains an ongoing scientific challenge for predicting adverse outcomes (Spinu et al., 2020; Zhou & Sahlin, 2026). To derive benchmark dose (BMD) thresholds from qAOPs built from in vitro assays, these frameworks must model the non-linear relationships in biological networks. In order to make reliable predictions and to establish thresholds for the adverse outcomes, qAOP models must estimate the generalization error by quantification of the uncertainty bounds and correctly propagate non-linear variance.

Bayesian networks (BNs) are a promising approach for quantifying AOPs due to their capability to estimate model uncertainties within a network and to produce probabilistic predictions (Moe et al., 2021; Spinu et al., 2020; Tebby et al., 2022; Zhou & Sahlin, 2026). To date, however, the application of Bayesian networks is still subject to the following limitations: Initially, qAOPs (Moe et al., 2021) using Bayesian regression and converging pathways employed discretization of continuous biological data into categorical classes, which impairs predictive power and precision (Cao et al., 2023). Moreover, their approach relied on randomization of the order of observations in each dose group of the assay to resolve the challenge that observations of different key event relationships are only coupled by dose. The use of continuous linear Gaussian networks was proposed by Ito et al. (2024), which resolves the discretization problem for converging pathways. By design, linear models cannot capture non-linear biological properties like low-dose activation and high-dose saturation effects, which can result in low predictive power (Durnik et al., 2026). Continuous, non-linear Bayesian inference without the need for discretization was proposed by Tebby et al. (2022), but their approach was limited to single sequential pathways. The result is that the individual pathways in AOP networks can often only be analyzed as separate Bayesian networks (Tebby et al., 2022; Zhou & Sahlin, 2026), or only linear models are used for relationships that are actually non-linear (Cao et al., 2023; Durnik et al., 2026; Ito et al., 2024). Recently, Zhou and Sahlin (2026) have overcome some of these limitations by using latent variables to estimate Bayesian regression models between key event relationships without bias, and proposed probabilistic Bayesian leave-one-out cross-validation (LOO-CV) for uncertainty estimation to evaluate the model fit for multiple compounds.

However, two important methodological gaps remain: No existing approach offers a mathematically viable, transparent method for merging and weighting multiple converging, non-linear pathways in a Bayesian network. When multiple signaling pathways, such as those of the retinoic acid receptor (RAR), pregnane X receptor (PXR), farnesoid X receptor (FXR), and peroxisome proliferator-activated receptor gamma (PPARγ), converge on a single adverse outcome, researchers lack a robust statistical method to isolate the most important biological drivers from redundant pathways. Moreover, no accurate, generally applicable framework for uncertainty estimation suitable for smaller datasets with missing data has yet been tested for continuous Bayesian AOP networks.

Here, we present a probabilistically weighted Bayesian qAOP Ensemble (Bayesian qAOPE) that combines converging molecular signaling pathways within an AOP network using exact Bayesian LOO-CV stacking. To mathematically model the key event relationships that are related only in terms of dose, we use latent variables (Zhou & Sahlin, 2026). Adapting the spline approach of Cao et al. (2023), we used heteroscedastic, nonlinear dose-response models and heteroscedastic thin-plate splines to model potential activations in the low-dose range and potential saturation effects in the high-dose range. As a result, the complete posterior predictive distributions are propagated through the network, with epistemic parameter uncertainty and aleatory biological variance (Sahlin et al., 2021) incorporated into the predictions. The application of our Bayesian framework demonstrates that the Bayesian qAOPE is suitable as a macroscopic prediction filter for triglyceride accumulation (TGA) in an in vitro AdipoRed assay. The Bayesian qAOPE identified the network’s RAR signaling pathway as the dominant driver (83.9% weight) of the adverse outcome, while simultaneously consolidating redundant paths of the network. With a 95.8% out-of-sample predictive coverage, the Bayesian qAOPE exhibits an exceptionally high predictive capacity. The Bayesian qAOPE framework offers a blueprint for Bayesian-based probabilistic predictions within qAOPs and the estimation of a network-derived benchmark dose (BMD) value with minimal uncertainty.

## 2. Materials and Methods

### 2.1 Data

The quantitative adverse outcome pathway (qAOP) for liver steatosis was based on dose response data from Knebel et al. (2019), where cell lines were treated with the plant protection product (PPP) propiconazole. Dose response data for MIEs were produced by the dual-luciferase reporter (DLR) assay by measuring the ability of propiconazole to affect the transcriptional activity of different nuclear receptors in HepG2 cells. Dose response data on triglyceride accumulation (TGA) were produced by the AdipoRed assay using HepaRG cells.

The dataset will be provided in the supplementary material. The statistical computing environment R (R Core Team, 2026) was used for all data analyses and modeling described below. The data were processed using the R packages dplyr (Wickham et al., 2026) and tidyr (Wickham et al., 2025).

The experimental datasets used in this analysis included independent, multi-test toxicity assessments targeting specific key events: Molecular initiating events (MIEs) (PXR, FXR, RAR, PPARγ) and the adverse outcome of triglyceride accumulation (TGA).

The dose-response relationships for the molecular initiating events (MIEs) spanned propiconazole concentrations from 0 to 40 µM, while the triglyceride accumulation (TGA) response was measured from 0 to 20 µM. The measurements for the control group for all key events were normalized to 1. We assumed log-normal distributions for all endpoints. Thus, all response data were log-transformed prior to modeling to avoid negative values on the original scale (Slob, 2002; Tebby et al., 2022).

### 2.2 Adverse Outcome Pathway (AOP)

An adverse outcome pathway (AOP) network for liver steatosis was constructed based on the AOP proposed by Knebel et al. (2019) (see also (Vinken, 2016)) and adapted to the available data (Figure 1). The molecular initiating events (MIEs) were PXR, FXR, and RAR. It is assumed that all MIEs act as agonists and are upregulated by increasing doses of the steatosis-inducing substance (Knebel et al., 2018; Marx-Stoelting et al., 2020). The activation of PPARγ was modeled as a downstream Key Event of PXR, establishing a key event relationship (Foran et al., 2019; Mellor et al., 2016; Moe et al., 2021). All MIEs were linked to the adverse outcome, triglyceride accumulation (TGA) (see Figure 1).

**Figure 1:**
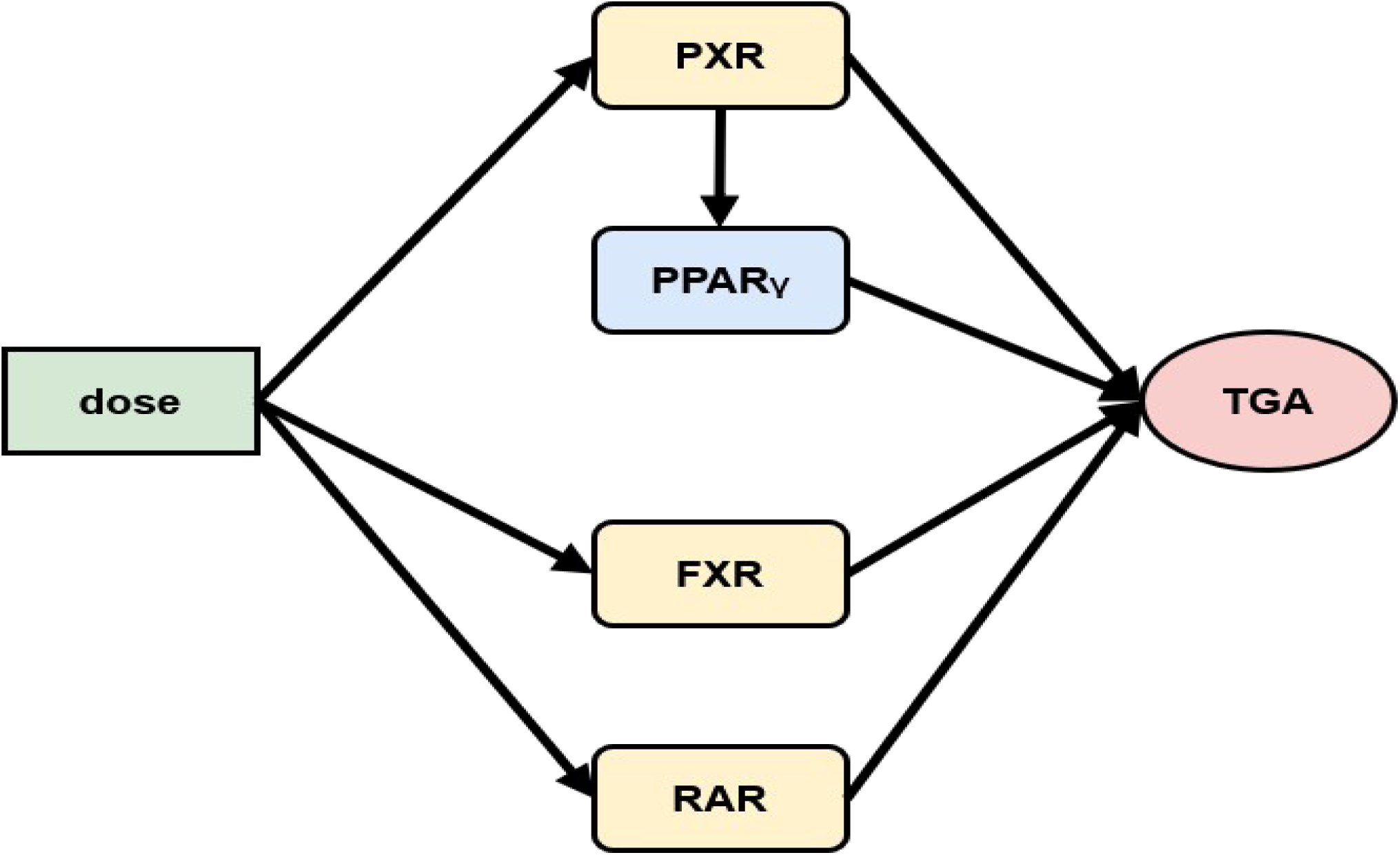
Quantitative adverse outcome pathway (qAOP) for the adverse outcome triglyceride accumulation (TGA). Dose is the potentially adverse substance. Molecular initiating events (MIEs) directly linked to dose are PXR, FXR, and RAR. The downstream Key Event PPARγ is shown in light blue. Structure of the AOP adapted according to available data from Knebel et al. (2019) and Vinken et al. (2015).

### 2.3 Empirical Benchmark Dose Derivation of TGA

Following Luckert et al. (2018), a biologically adverse outcome in the AdipoRed assay is defined at a benchmark response (BMR) of a 50% extra risk. Graphical analysis of the dose-response data showed evidence for signal variance increasing at higher doses (heteroscedasticity). To conduct this analysis and explicitly model the heteroscedasticity, we implemented a custom Bayesian distributional regression utilizing the R package brms (Bürkner, 2017, 2018, 2021). The dose-response relationship was modeled using a mathematically simplified three-parameter exponential model (with the background response fixed to 1) (Slob, 2002). This exponential model was selected because the dose-response data did not show an upper asymptote when visually inspecting the tested data. To address the heteroscedasticity, the residual standard deviation was modeled as a continuous linear function dependent on the dose.

Instead of uniform prior distributions, which are known to lead to boundary-induced bias (Wheeler, 2023), weakly informative priors were employed for all model parameters of all Bayesian regression models in this study, based on plausible biological responses (Lemoine, 2019). As the controls were normalized to one and to ensure a positive outcome for downstream key events, an informative prior was employed for the lower asymptotes (Lemoine, 2019). The weakly informative priors for regularization of all models in this study will be stated in the supplementary material.

Posterior distributions of the expected mean response were generated for a series of continuous doses to estimate the benchmark dose metrics. The dose at which the posterior median of the mean response curve intersected the BMR threshold was defined as the empirical BMD. The doses at which the 95% upper limit and the 5% lower limit of the mean response curve crossed the activation threshold were used to estimate the BMDL and the BMDU of the 90% BMD credible interval (CI).

### 2.4 Bayesian Dose-Response Modeling of Molecular Initiating Events (MIEs)

The dose-response relationships for FXR, RAR, and PPARγ were modeled by a four-parameter log-logistic function (Slob, 2002). By visual inspection, the PXR dose-response relationship did not show an upper asymptote in the tested data and thus was modeled by a three-parameter exponential model (Slob, 2002). Models were estimated by a No-U-Turn Sampler (NUTS) Hamiltonian Monte Carlo (HMC) sampler implemented in Stan (Carpenter et al., 2017) with the brms package in R (Bürkner, 2017, 2018, 2021). Heteroscedasticity was modeled as linear function dependent on the doses. Weakly informative priors were set for all model parameters as described in section 2.3.

Sampling was performed with 20,000 iterations and four chains for every model. The target average proposal acceptance probability, δ, was increased to 0.95 to prevent divergent transitions and increase the exploration of the posterior distribution. Sampling yielded posterior probability distributions for all model parameters. Convergence of all model parameters was checked by the potential scale reduction factor (*R̂*), bulk and tail effective sample sizes.

### 2.5 Distributional Non-Linear Modeling of the Downstream Events and the Adverse Outcome

The assays used to generate the dataset were independent of one another, resulting in a dataset where measurements between each dose group are only related with respect to the dose. The application of regression under the assumption that each row represents paired measurements between the key events violates the assumption of independent observations. To account for this heterogeneity in the trial series, a two-stage Bayesian framework (Tebby et al., 2022; Zhou & Sahlin, 2026) was used, which we extended by modeling heteroscedastic variance. First, the underlying, latent biological pathway was estimated by modeling each primary ‘dose-response’ relationship (see Section 2.4). Then standardized posterior expectations of the dose-response models were calculated, resulting in independent predictors for downstream key events. This step was repeated for all key event relationships.

To model the key event relationships to the adverse outcome (TGA), we used “thin-plate” type splines (k=4) (generalized additive models (GAM)) to evaluate the response-response relationships between key events (Cao et al., 2023; Wood, 2017), expanded here to heteroscedastic Bayesian models. The usage of splines accounts for the combination of threshold at the low doses and saturation effects at the top doses. Because of the limited sample size, a Gaussian prior distribution was applied to constrain model parameters and the variance penalty of the curve in order to prevent overfitting (Bürkner, 2017; Wood, 2017). Weakly informative priors were set for all model parameters as described in section 2.3.

The thin-plate splines were also compared to linear and heteroscedastic linear models for each key event relationship by Bayesian leave-one-out cross-validation (LOO-CV) with Pareto smoothed importance sampling (PSIS), as recently proposed for model selection in qAOPs (Zhou & Sahlin, 2026). If the Pareto k threshold was exceeded, posterior refitting was employed with the R package loo (Vehtari et al., 2024, 2025). The most parsimonious model was selected, if no meaningful difference could be found by the expected log predictive density criterion, Δ *ELPD* < *SE*, (Sivula et al., 2020; Vehtari et al., 2017). We accounted for the heteroscedasticity by using a distribution-based approach and quantifying it as a linear function of the predictor. The modeling was performed with the R package brms (Bürkner, 2017, 2018, 2021) with 20,000 iterations and four chains, a target average proposal acceptance probability of 0.999 and a maximum tree depth of 15.

### 2.6 Out-of-Sample Validation of Dose-Response and Response-Response models

The Bayesian LOO-CV with PSIS procedure also yielded an out-of-sample estimate of the predictive performance of the models. The Median and the 95% credible intervals (CI) of the Bayesian LOO-R^2^ (Gelman et al., 2019) were estimated using the R package loo (Vehtari et al., 2025) to estimate the out-of-sample predictive performance by the explained variance of the downstream connections from the key events PXR, FXR, RAR and PPARγ to the adverse outcome TGA.

### 2.7 Benchmark Dose Estimation and Propagation of Epistemic Uncertainty

A grid of doses was propagated through the stacked Bayesian qAOP Ensemble to sample draws from the expected value (mean response) of the posterior predictive distribution. Based on these draws, a BMD calculation was performed for an activation threshold of a 1.5-fold change above the estimated baseline. The BMD of the Bayesian qAOP network was estimated as the dose where the posterior probability of exceeding the activation threshold was 50%. A 90% credible interval was established in the same way by calculating the benchmark dose lower bound (BMDL) and benchmark dose upper bound (BMDU) as the doses where the posterior probabilities of exceeding the activation threshold were 0.05 and 0.95.

The probability of exceeding the threshold was plotted against dose as predictors to graphically compare the BMD and credible interval of the Bayesian qAOP network with the empirical BMD (Figure 6).

### 2.8 Bayesian qAOP Ensemble

The pathways of the qAOP network were modeled independently and each pathway can be treated as a separate Bayesian network. For modeling the convergence of independent pathways going through the nodes PXR, FXR, RAR and PPARγ to the same node, in this case to the adverse outcome node TGA, a Stacked Bayesian Ensemble was employed (Yao et al., 2018): predictions for the adverse outcome were sampled from the posteriors and then weighted by their Bayesian LOO-CV stacking weights.

Stacking allows the combination of pathways with strongly correlated outcomes, as all pathways in the network model agonistic (increasing) dose-response relationships. The stacking weights were also employed to assess the relative importance of each pathway, weighting down pathways that were redundant in their explanation of the adverse outcome (Yao et al., 2018).

### 2.9 Forward Sampling and Posterior Predictive Analysis

A posterior predictive check of the Bayesian qAOP Ensemble was performed by forward sampling (Henrion, 1988; Koller & Friedman, 2009; Scutari & Denis, 2021), based on a dose grid from dose 0 to dose 20 (stepsize = 0.01) and on the tested doses (0, 0.625, 1.25, 2.5, 5, 10, 20) (Figure 7), each with 4000 posterior predictive draws per node. Forward sampling through the Bayesian qAOP Ensemble propagates parameter uncertainty and biological noise through the qAOP network (Gelman et al., 2013). The 4000 posterior predictive draws were parallelized with the R package future.apply (Bengtsson, 2021) and the progress visualized with the R package progressr (Bengtsson, 2026).

### 2.10 Estimation of the Generalization Error by Bayesian Leave-One-Out Cross Validation

To estimate the generalization error including the parameter uncertainty and the uncertainty due to biological variance (epistemic and aleatoric uncertainty) (Sahlin et al., 2021), an exact Bayesian LOO-CV algorithm (Vehtari et al., 2017) was employed: Each observation of the adverse outcome TGA was removed from the dataset as a held-out test set. All parts of the Bayesian qAOP Ensemble dependent on TGA were completely re-estimated on the remaining data (train set) as described above. Then the dose value of the held-out test value was propagated through the network by forward sampling as described in section 2.8.

The results of the predictions were evaluated for each held-out test value by calculating the 50% and 95% predictive coverage intervals to check that the empirical observations were correctly described by the model (Figure 9). Moreover, the empirical cumulative distribution function (ECDF) of the Probability Integral Transform (LOO-PIT) was estimated in order to verify that the predictive distributions were uniform (Figure 8) (Dawid, 1984; Gabry et al., 2019). For the posterior predictive model visualizations the R package bayesplot (Gabry et al., 2019) was used.

## 3. Results

### 3.1 Fits of Dose-Response and Response-Response Models

For the MIES (dose to PXR, RAR and FXR), dose-response models were fitted as described in Section 2.4 (Figure 2).

**Figure 2:**
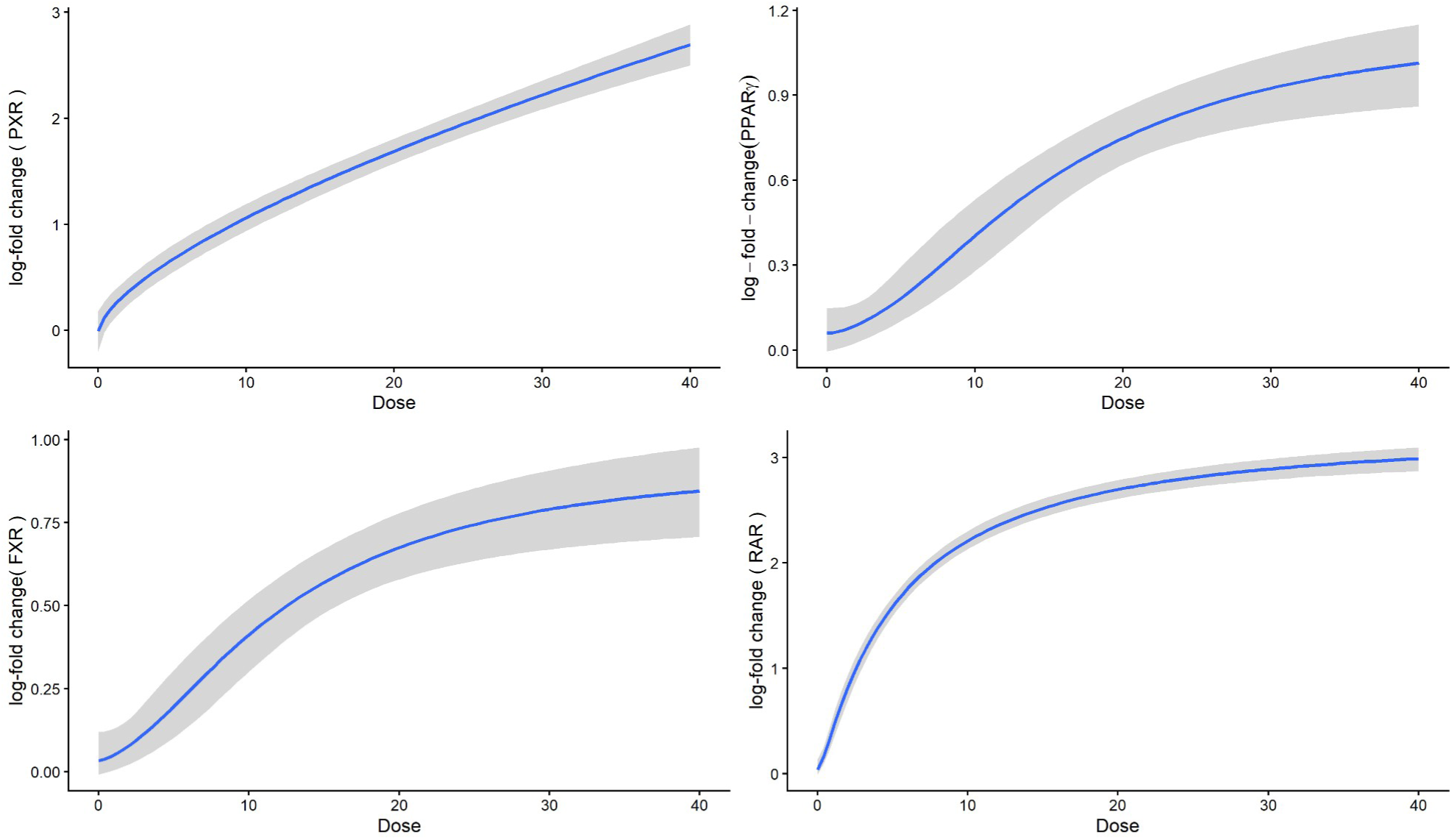
Non-linear dose-response models of the molecular initiating events (MIEs) PXR, FXR, PPARγ, and RAR. The dark blue line represents the posterior median model fit, the 95% credible interval is depicted by the shaded region. A four-parameter log-logistic model (with the intercept fixed to one on the original scale) was fitted for PPARγ, FXR, and RAR and a three-parameter exponential model was employed for PXR.

The key event relationships of PXR, RAR and (cascaded) PPARγ to the adverse outcome, TGA, were modeled by non-linear thin-plate spline models (Figure 3b), because LOO-CV showed a clearly better fit of the thin-plate spline models when compared to the linear heteroscedastic models (PXR (Δ *ELPD*=4.7 *, SE* 1.9), PPARγ (Δ *ELPD*=4.6 *, SE*=1.9), and RAR (Δ *ELPD*=4.1 *, SE*=2.3)). The choice of the spline models was corroborated by visually identified high dose saturation and low dose activation effects.

**Figure 3a:**
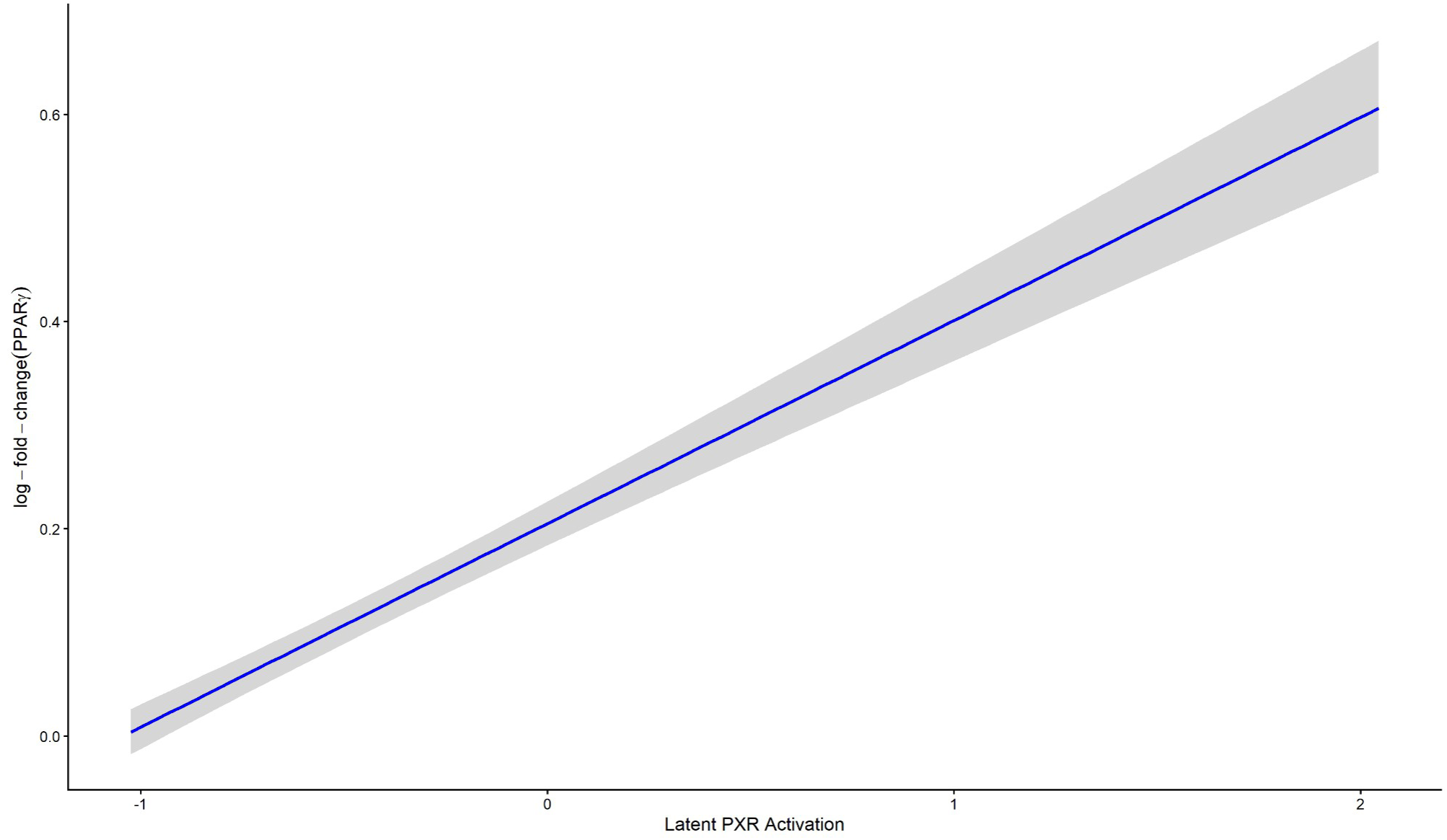
Key event relationship from PXR to PPARγ. The dark blue line depicts the posterior median fit and the gray shaded region the 95% credible interval (log-scale) of the fitted heteroscedastic linear model.

**Figure 3b:**
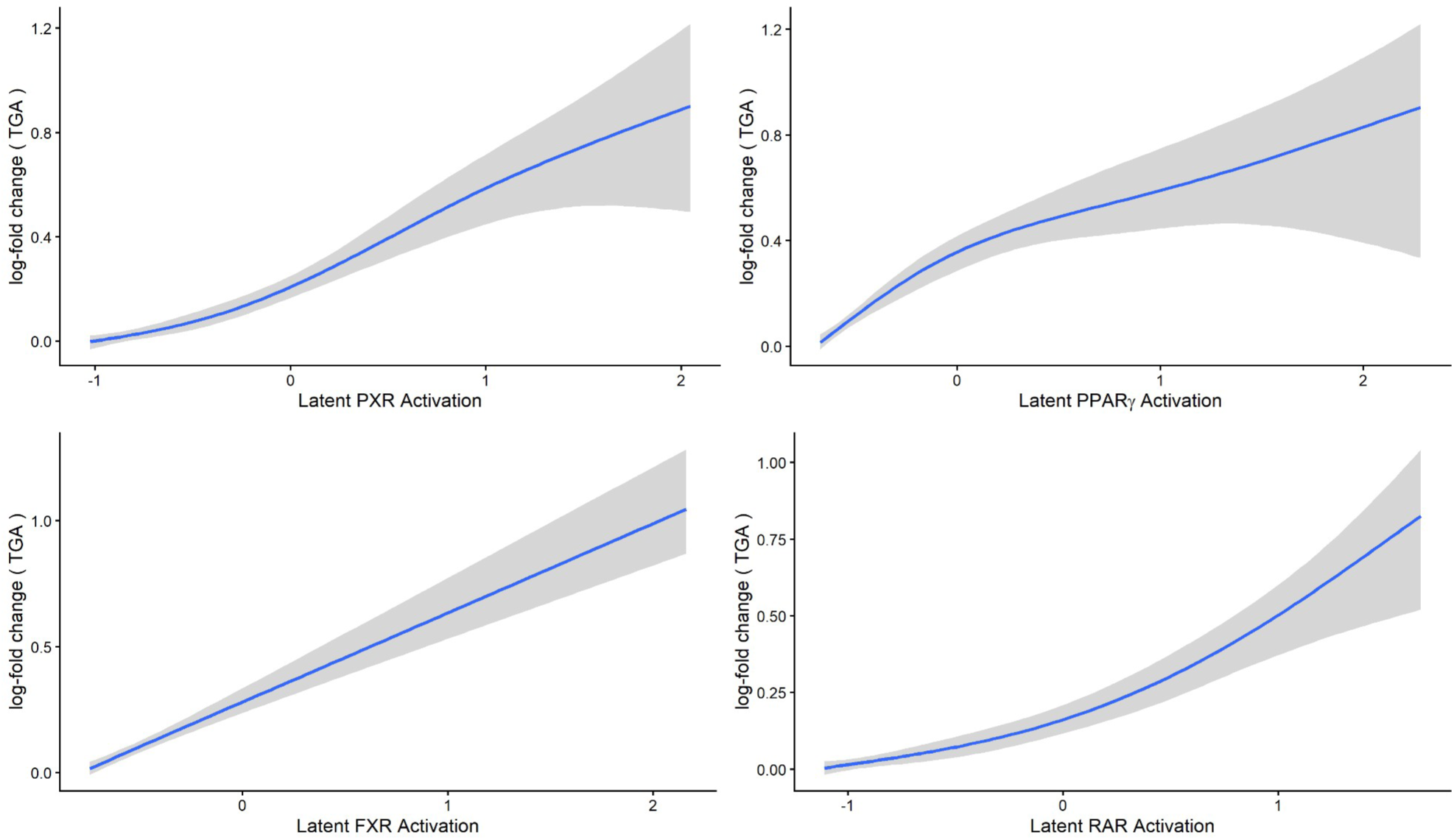
Response-response models mapping the key event relationships from the continuous latent predictors (PXR, FXR, RAR, and PPARγ) to the adverse outcome (TGA) (log-scale). The dark blue line represents the posterior median fit. The shaded area marks the 95% credible interval. Relationships were estimated by penalized heteroscedastic thin-plate splines (k=4) except for the key event relationship from FXR to TGA, where a heteroscedastic linear model was fitted.

For the key event relationship of FXR to TGA, LOO-CV did not show a meaningful difference of the spline model when compared to the heteroscedastic linear model (Δ *ELPD*=0.9 *, SE*=1.2), so the more parsimonious heteroscedastic linear model was chosen. Fitting the spline model to the key event relationship of PXR to PPARγ led to 160 divergent transitions and the difference between the spline and the heteroscedastic model measured by the LOO-CV was not meaningful either (Δ *ELPD*=0.6 *, SE*=0.6), so the heteroscedastic linear model was selected (Figure 3a).

All models were successfully estimated (*R̂* ≤1.01, bulk and tail effective sample size (ESS) of several thousand iterations) and graphically checked for plausibility (Figures 2, 3a and 3b). Standardized residuals of all fitted models corroborated the successful model estimation being centered around zero with nearly all observations falling into the [-2, 2] interval.

The fit of the response-response models between the key events and the adverse outcome, TGA, (epistemic uncertainty) was also evaluated by the explained out-of-sample variance (LOO-R^2^). The response-response-model from FXR to TGA had the highest LOO-R^2^ (90.7% [95% CI: 77.6% - 97.0%]), followed by the key event relationship from PPARγ to TGA (90.3% [95% CI: 81.8% - 93.6%]), PXR to TGA 90.2% [95% CI: 82.0% - 93.5%], and RAR to TGA (87.4% [95% CI: 79.3% - 92.9%]) (Figure 4).

**Figure 4:**
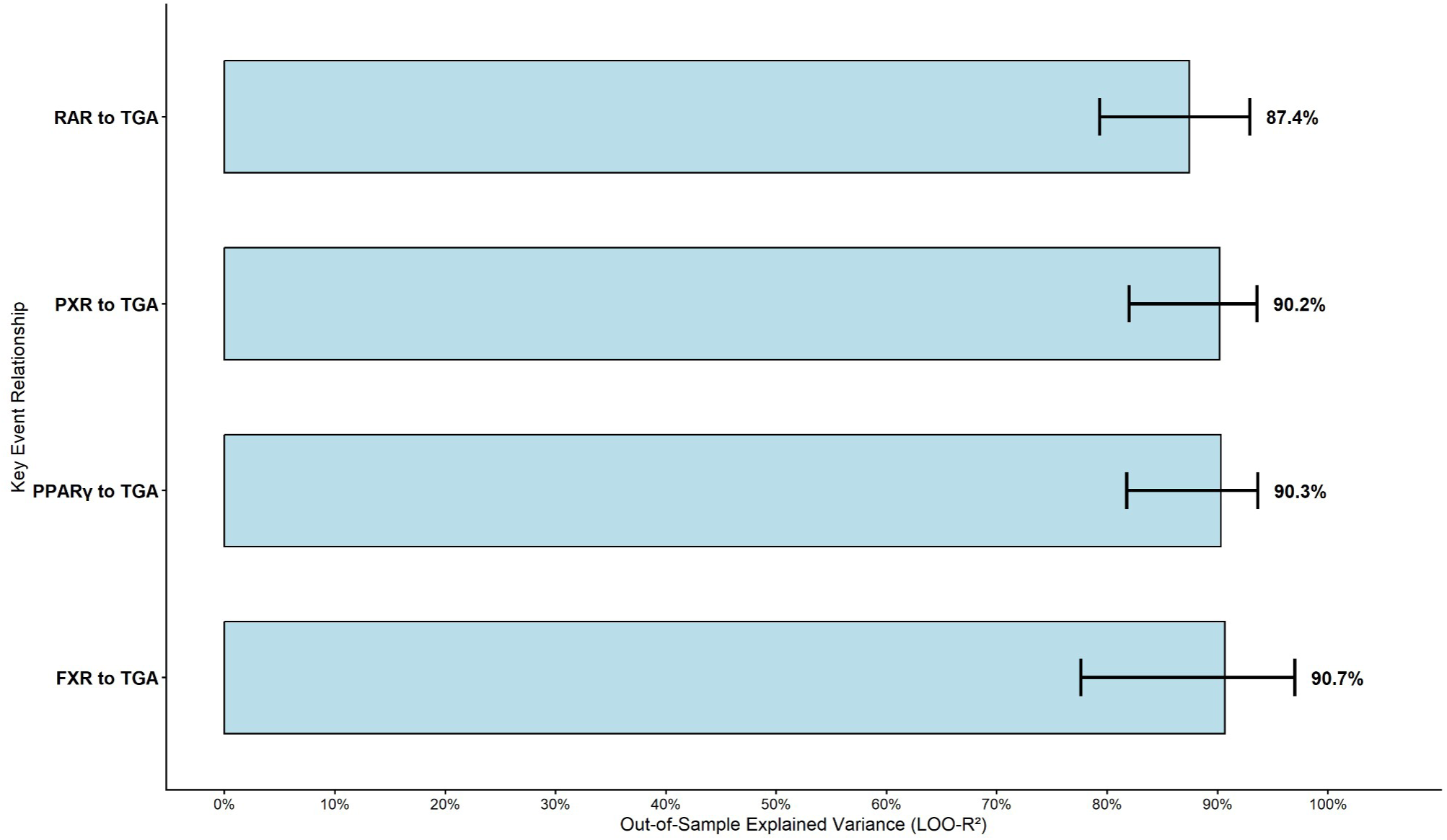
Out-of-sample explained variance (LOO-R²) of the key event relationships of the MIEs to the adverse outcome, TGA, within the Bayesian qAOP Ensemble. Horizontal bars show the estimated LOO-R² values and the black error bars the 95% credible intervals. The median values are stated to the right of the 95% credible intervals.

All plots were generated using the R packages ggplot2 (Wickham, 2016) and patchwork (Pedersen, 2025) and bayesplot (Gabry et al., 2019).

### 3.2 Contribution of Pathways to the Bayesian qAOP Ensemble

The contribution of the pathways to the Bayesian qAOPE was governed by the Bayesian stacking weights. Analysis of the Bayesian stacking weights governing the contribution of the pathways to the Bayesian qAOP Ensemble showed that the clearly highest weight (83.9%) was assigned to the pathway via RAR with a minor contribution of the pathway connecting PXR to TGA (16%). The pathways via FXR and PPARγ had negligible effect on the final outcome (0.0% and 0.1%) (Figure 5).

**Figure 5:**
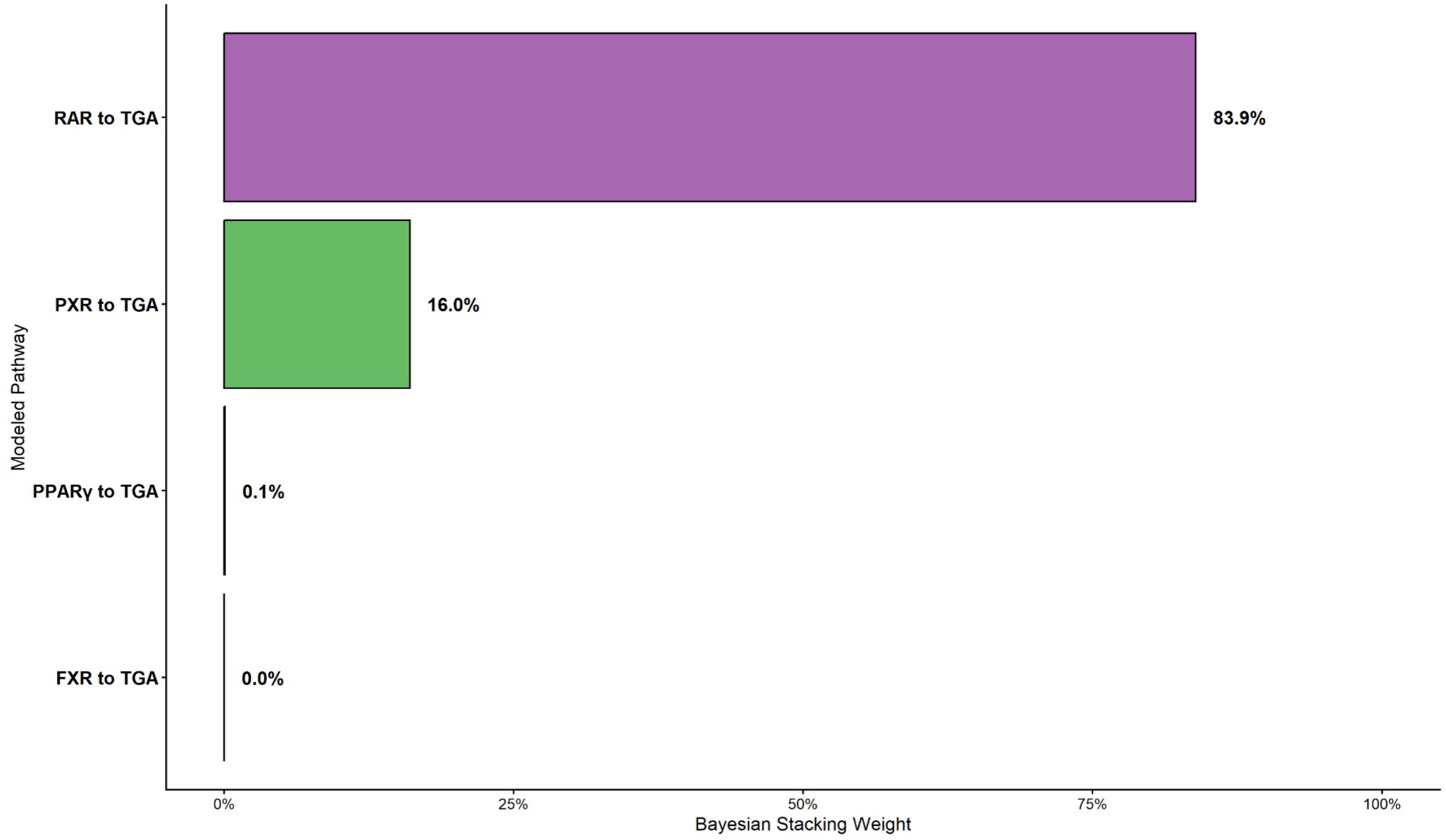
Relative predictive weight of pathways within the Bayesian qAOP Ensemble (Bayesian qAOPE). Bars show the Bayesian Stacking Weights of Leave-One-Out (LOO) Expected Log Predictive Densities (ELPD). The Bayesian Stacking weights determine the weight of each pathway to the adverse outcome TGA for the estimated Bayesian qAOPE.

### 3.3 Comparison of Empirical and Probabilistic Benchmark Dose Estimation

The three-parameter exponential model was successfully fit to estimate the empirical relationship between dose and TGA as described in section 2.3 and 2.4 (*R̂* ≤1.01, bulk and effective sample size of several thousand iterations, and residuals centered around zero in the interval [-2,2] (Figure 6A).

**Figure 6:**
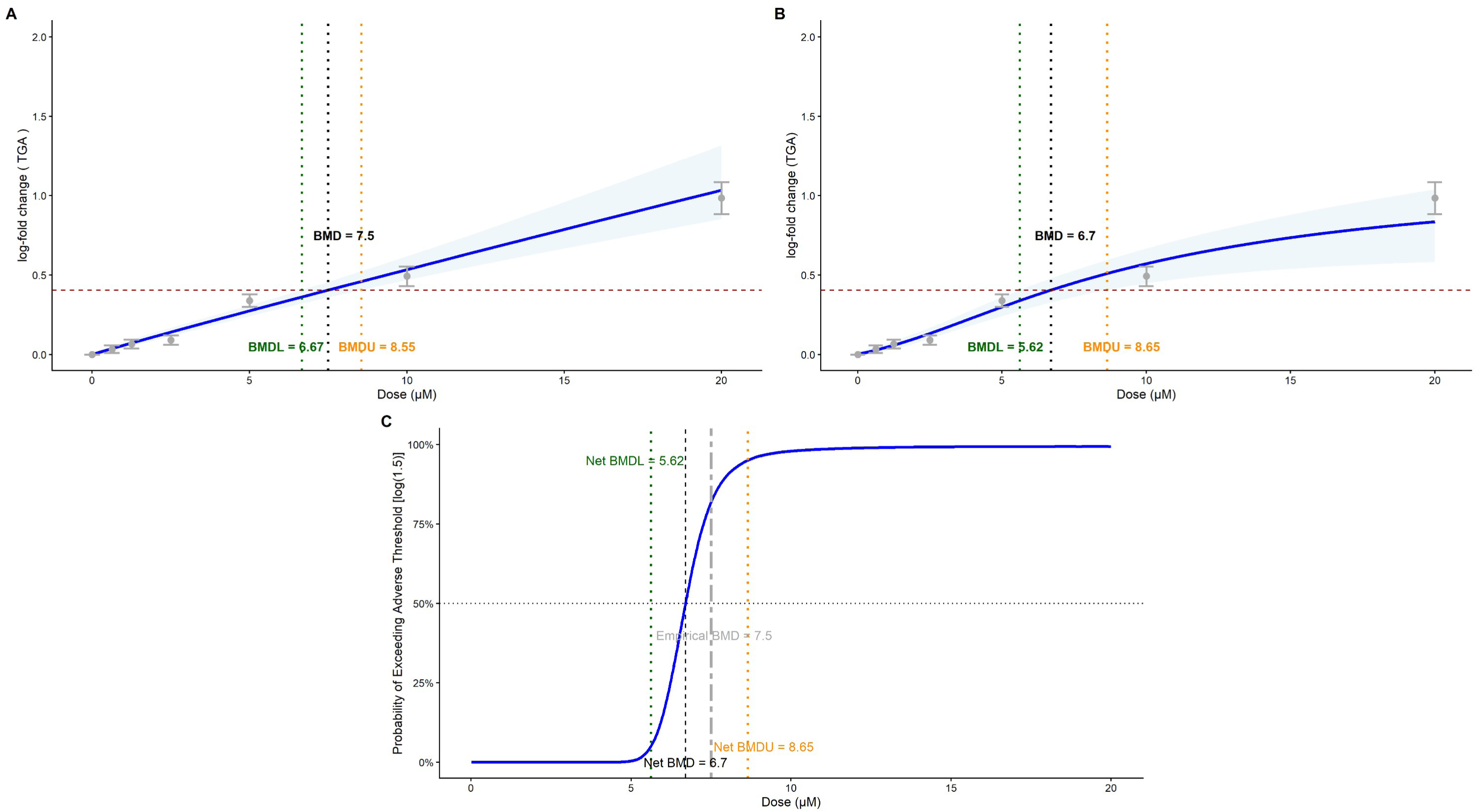
Probabilistic Benchmark Dose (BMD) derivation for the Bayesian qAOP Ensemble (Bayesian qAOPE). **(A)** (BMD analysis of empirical measurements of TGA after treatment with propiconazole) The empirical three-parameter exponential model led to a median BMD of 7.5 µM (black dashed line) and a 90% credible interval of [6.67, 8.55]. The dark blue line depicts the median posterior expectation, with the 95% credible interval shaded in light blue. The empirical measurements at each dose of propiconazole are depicted with grey dots, with the error bars indicating +/-1 standard error of the mean (SEM). **(B)** (Adverse outcome TGA simulated by Bayesian qAOPE and BMD analysis)The dose-response curve simulated by the Bayesian qAOPE leads to a BMD estimate of 6.7 µM and a 90% credible interval of [5.62, 8.65]. The dark blue line represents the median posterior expectation, with the 95% credible interval shaded in light blue. The empirical measurements at each dose are depicted with grey dots, with the error bars indicating +/-1 standard error of the mean (SEM). **(C)** The probability of exceeding the BMD derived by the Bayesian qAOPE (black dashed line) is depicted by the blue curve. The BMD estimated by the Bayesian qAOP Ensemble and its 90% credible interval are shown by the black, green and orange dotted lines.

The estimated empirical Benchmark Dose (BMD) value for a 50% extra risk was 7.5 µM, with a 90% BMDL of 6.67 µM and a 90% BMDU of 8.55 µM. The resulting BMDU/BMDL ratio of 1.3 signals a low uncertainty in the model fit.

The Bayesian qAOP Ensemble (Bayesian qAOPE) returned a simulated BMD of 6.7 µM, with a 90% BMDL of 5.62 µM and 90% BMDU of 8.65 µM. The BMDU/BMDL ratio of the Bayesian qAOPE was 1.5, comparable to the empirical model. Both the empirical BMD and the BMD estimated via the Bayesian qAOPE fell within their respective credible intervals.

### 3.4 Posterior Predictive Check of Bayesian qAOP Ensemble

Posterior predictive simulation of the Bayesian qAOPE by 4000 MCMC forward sampling draws at the tested doses of TGA led to 100% of the observations (24 out of 24) falling within the 95% posterior predictive interval (PPI), and 58.3% of the observations (14 out of 24) into the 50% PPI. The results of the posterior predictive check via forward sampling is shown for the dose grid in Figure 7. It can be seen in both figures that the non-linear, heteroscedastic increase of variance at higher doses is successfully captured by the Bayesian qAOPE. There is no variance explosion at either the tested or the interpolated doses (Figure 7).

**Figure 7:**
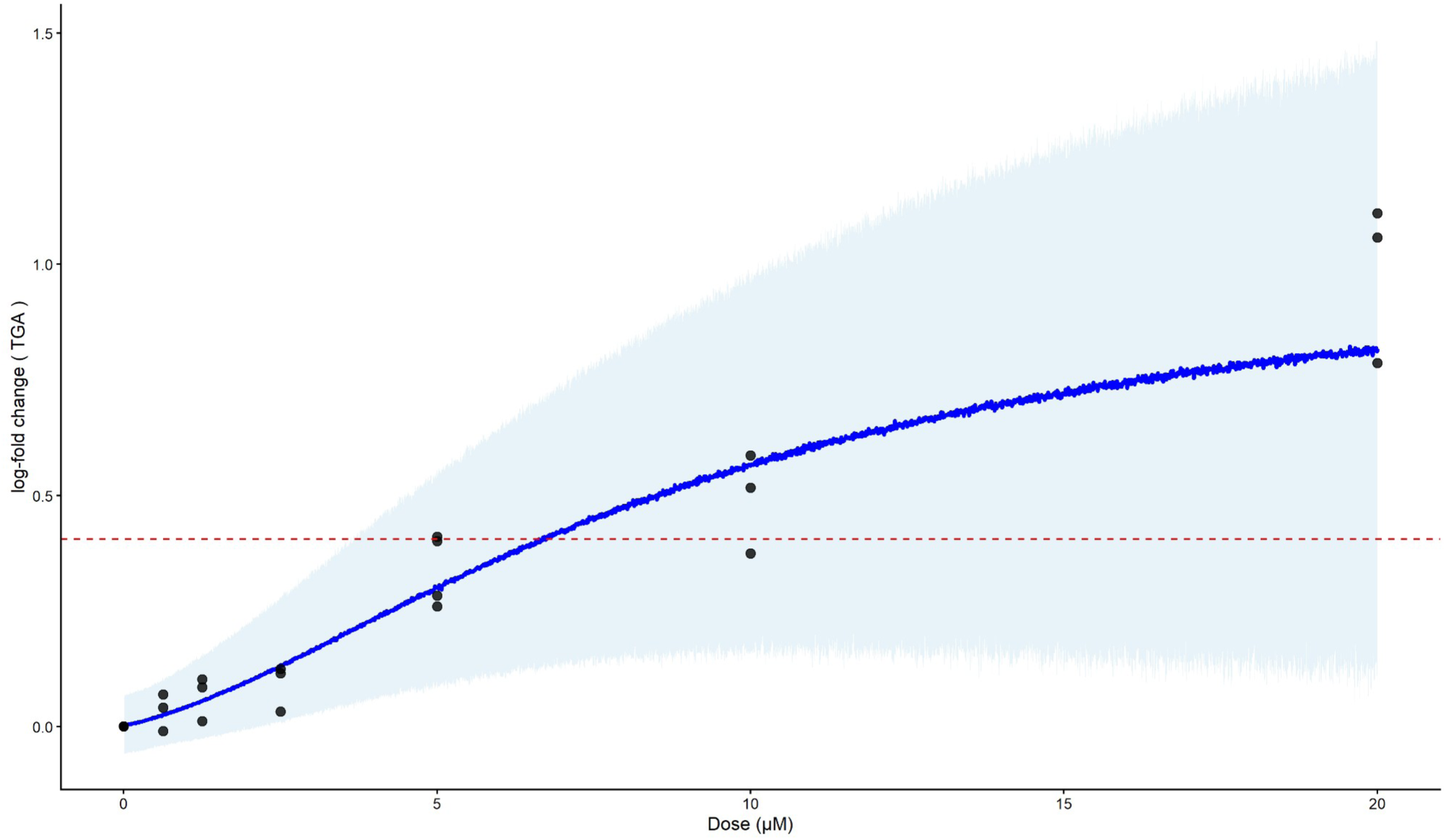
Posterior predictive check of the Bayesian qAOP Ensemble (Bayesian qAOPE). A grid of 2000 nodes (stepsize = 0.01) from dose 0 to 20 µM of propiconazole was propagated through the Bayesian qAOPE by forward sampling for 4000 MCMC draws. The median of the simulated values is depicted by the dark blue line and the 95% posterior prediction interval (PI) by the light blue shaded area. Measurements of triglyceride accumulation (TGA) at the doses are plotted as black dots. The activation threshold of log(1.5) (50% extra risk on the original scale) is marked by the red dashed line. It can be seen that the variance stays bounded at the measured doses and the interpolated area. The simulations capture the non-linear, heteroscedastic behaviour observed in the tested doses.

### 3.5 Bayesian Exact Leave-One-Out Cross-Validation Estimate of the Generalization Error

Estimation of the generalization error of the Bayesian qAOPE by exact Bayesian leave-one-out cross-validation (LOO-CV) yielded 95.8% of the true TGA measurements (23 out of 24) falling within the 95% predictive interval, and 54.2% (13 out of 24) of the true TGA measurements falling into the 50% predictive interval (Figure 9).

The calibration of the empirical cumulative distribution function (ECDF) was successfully checked by the leave-one-out probability integral transform (LOO-PIT) method over the complete probability space (Figure 8).

**Figure 8:**
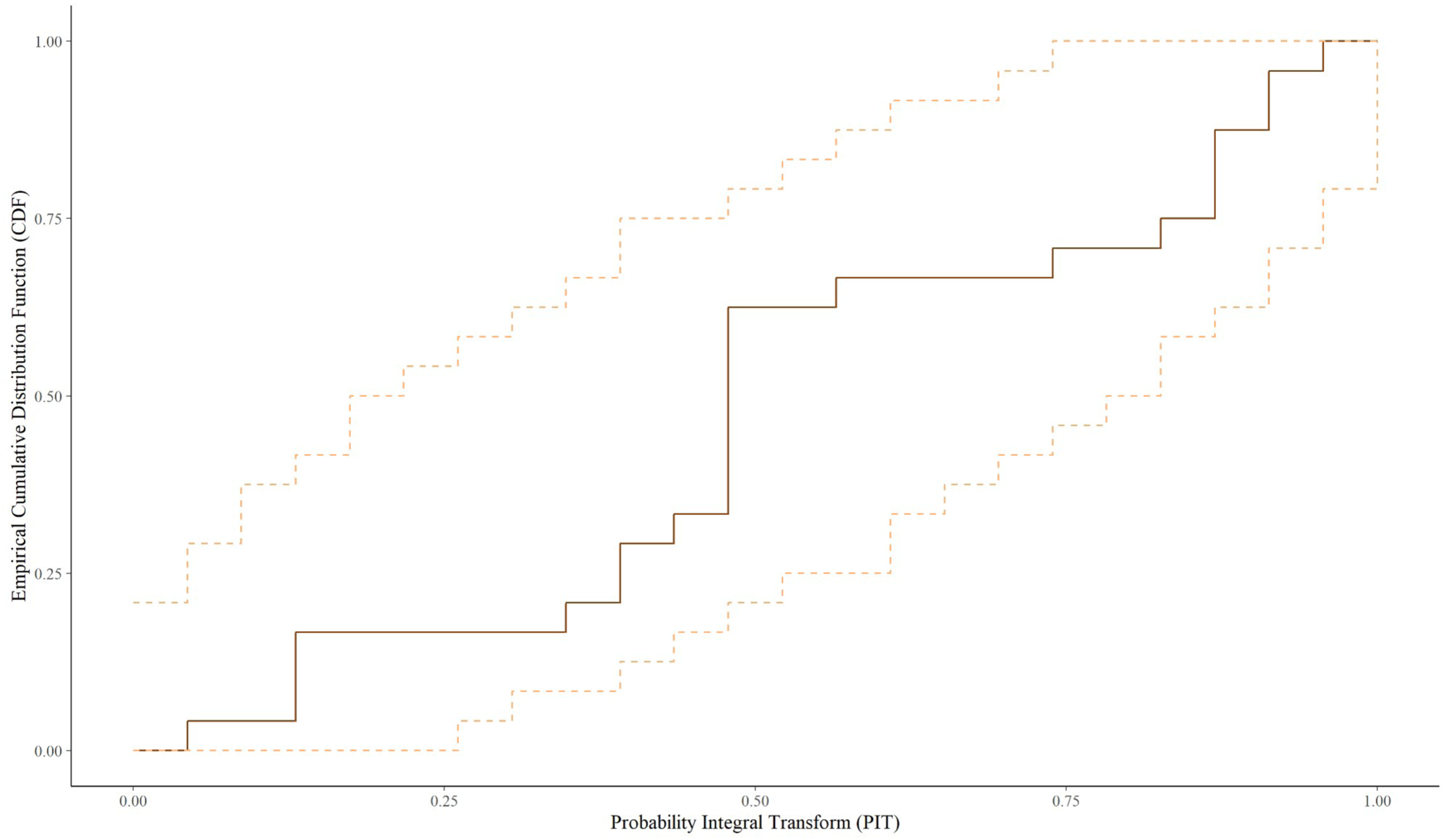
Exact leave-one-out probability integral transform (LOO-PIT). The empirical cumulative distribution function (ECDF) (solid orange line) is situated within the 95% confidence envelope of the uniform distribution over the whole range of probabilities (dashed orange lines). This shows the out-of-sample predictive calibration of the Bayesian qAOP Ensemble (Bayesian qAOPE).

**Figure 9:**
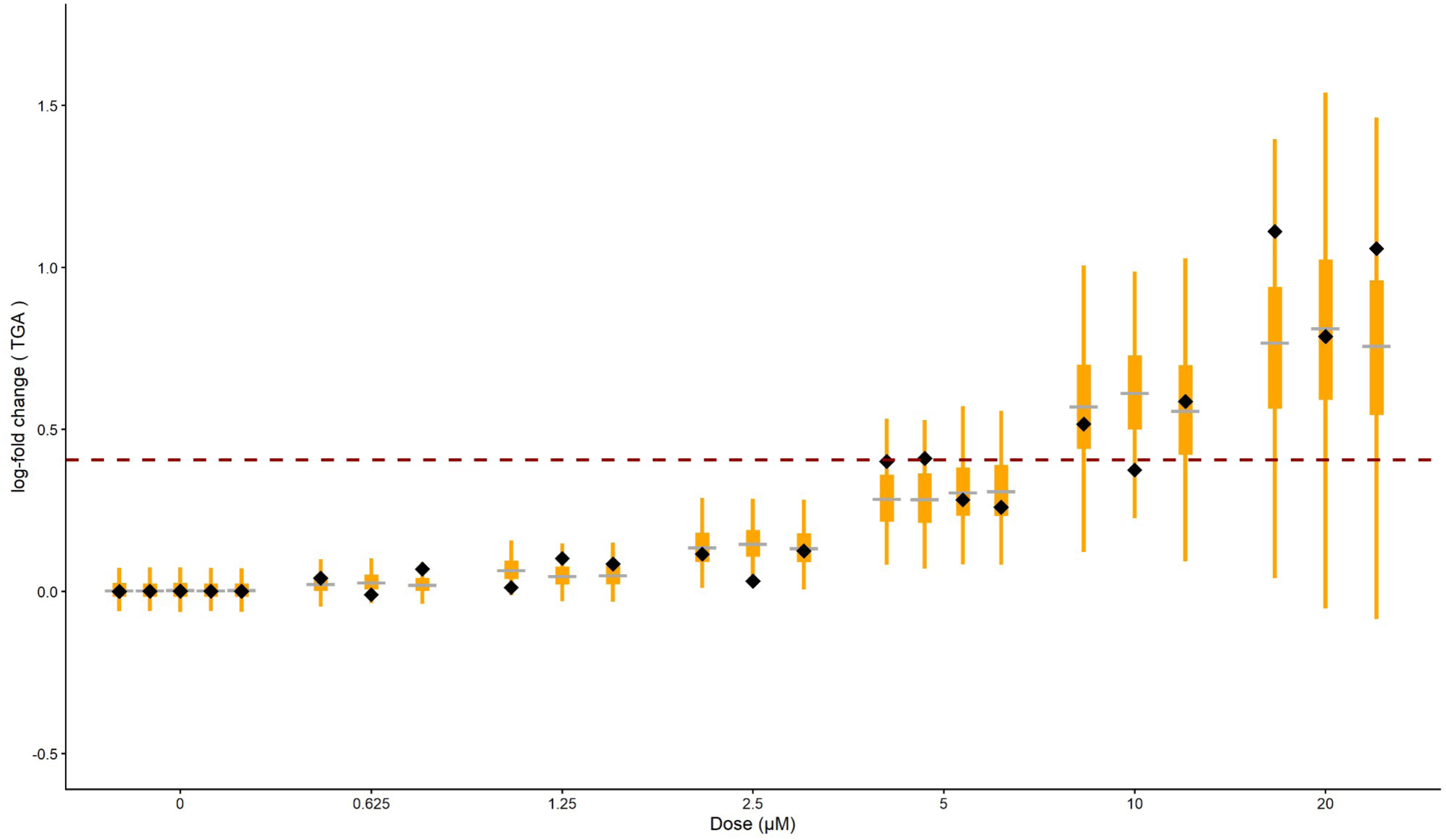
Estimates of the generalization error of the Bayesian qAOP Ensemble (Bayesian qAOPE) at the measured doses of triglyceride accumulation (TGA) for exact Bayesian Leave-one-out Cross Validation (LOO-CV). To estimate the generalization error, one measurement of TGA was left out as a test set before refitting the complete Bayesian qAOPE. The orange bars show the interquartile range and the 95% posterior predictive intervals of 4000 MCMC draws of forward sampling of the Bayesian qAOPE without the test set at the measured doses. The median of the simulation is in dark grey. The actual measurements of triglyceride accumulation (TGA) that were left out before the refit and the forward sampling simulation are depicted by the black diamonds. Of all measurements, 95.8% (23 out of 24) fell within the 95% posterior predictive interval, and 54.2% (13 out of 24 observations) fell within the 50% posterior predictive interval. The activation threshold at log(1.5) is depicted by the red dashed line.

## 4. Discussion

In this study, we developed a Bayesian quantitative adverse outcome pathway (AOP) Ensemble (Bayesian qAOPE) that yields highly accurate probabilistic predictions for the adverse outcome triglyceride accumulation. The Bayesian qAOPE integrates multiple converging pathways for deriving a biologically plausible continuous BMD value, which overcomes the dichotomous risk ratios proposed in (Zhou & Sahlin, 2026).

A critical issue in the application of Bayesian networks in toxicology is the risk of “overfitting” on small datasets. We used exact Bayesian leave-one-out cross-validation (LOO-CV) to show that the Bayesian qAOPE has exceptional out-of-sample predictive power, as it captures 95.8% of the empirical measurements within its 95% predictive interval. This generalization error demonstrates that the model correctly estimates the epistemic out-of-sample bounds, which is further confirmed by a BMDU/BMDL ratio of 1.5 signaling low uncertainty of the BMD estimate. Thus, the model’s out-of-sample bounds are governed by biological variance, rather than by overfitting.

Exact Bayesian LOO-CV displays a key property of the Bayesian qAOPE when compared to the in-sample posterior predictive check (PPC): While the in-sample PPC yielded an overly conservative coverage of 100% of the observations for the 95% posterior predictive interval (PPI) and 58.3% within the 50% PPI, the out-of-sample LOO-CV yielded nearly perfect coverage of 95.8% and 54.2%. In particular, the very high agreement with the nominal 50% coverage indicates that the high overall coverage of the Bayesian qAOPE is a consequence of the precise modeling of the central probability mass, and not due to artificially inflated uncertainty bounds. The out-of-sample metrics showing a slight decline when compared to the in-sample metrics corroborates that the heteroscedastic modeling of the paths learns the actual aleatory variance and transfers it to new observations.

Bayesian LOO-CV supported the conclusion that modeling response-response relationships using thin-plate splines led to vastly better fitting models, showing a clearly superior predictive density (Δ *ELPD*) for the splines when compared to linear models, as the splines were able to model the activation at low doses and saturation effects at high doses typical of receptor-binding compounds. By posterior forward sampling using our Bayesian qAOPE, the uncertainty of the model parameters (epistemic uncertainty) and the biological variance (aleatory uncertainty) (Sahlin et al., 2021) are propagated through the network. Forward sampling also yields prediction intervals that describe the entire distribution of biological variance, not just the confidence level for the mean. This results in high reliability (low uncertainty) of the derived BMD values (BMDU and BMDL).

The application of BNs to qAOPs with converging, non-linear pathways is still limited by the need for the discretization of continuous variables (Cao et al., 2023; Moe et al., 2021), the absence of paired observational data for linked key events (Durnik et al., 2026; Tebby et al., 2022) and the absence of a method for a true out-of-sample estimate of the generalization error for Bayesian qAOPs (Tebby et al., 2022; Zhou & Sahlin, 2026). Moreover, methods using cross-validation for model evaluation or pruning of continuous Bayesian networks (Durnik et al., 2026; Ito et al., 2024) are modeling non-linear pathways by linear Gaussian relationships, which can reduce predictive power.

We used latent variables to link observations from different assays only coupled by dose groups as proposed by Zhou and Sahlin (2026), which leads to an unbiased estimate of the empirical variance and mean. We found that this method was also suitable for a reliable model estimate on sparse datasets. Approximate LOO-CV has already been used for model selection of simple hierarchical paths in qAOPs involving multiple chemicals (Tebby et al., 2022; Zhou & Sahlin, 2026). To the best of our knowledge, this study is the first to propose Bayesian stacking based on exact LOO-CV as a method for estimating converging, nonlinear, and continuous qAOP cascades, thereby enabling the modeling, not only of simple, hierarchical pathways (Tebby et al., 2022; Zhou & Sahlin, 2026), but also of complex AOPs with converging pathways and non-linear relationships.

Bayesian stacking weights estimated from the expected logarithmic predictive density (ELPD) yield a data-driven weighting of the statistical influence of the AOP’s biological pathways on triglyceride accumulation. The clear dominance of the retinoic acid receptor (RAR) signaling pathway (83.9%) is consistent with its well-known role in lipid homeostasis. The direction of the RAR dose-response function is context-dependent. While in vivo models generally show that loss of RAR-alpha function drives hepatic steatosis (Li et al., 2024), in vitro models, especially under high-glucose conditions typical of cell culture assays, show that RAR activation paradoxically upregulates SREBP-1 and thus increases intracellular triglyceride accumulation (Blaner, 2019). This context-dependent metabolic switch is consistent with the upward trend of the RAR-TGA response-response function and the biological functioning of the in vitro AdipoRed assay. The pregnane X receptor (PXR) signaling pathway had a secondary but still measurable influence (16.0%), which can be explained by its known role as a sensor for exogenous substances and as a driver for hepatic lipid modulation (Li et al., 2024; Wang et al., 2023). Knebel et al. (2019) found that PXR plays an essential role in propiconazole mediated lipid accumulation. This is, however, not in contrast to our findings as the present models attributes an important role to PXR and PXR is also interacting with RAR. The minimal weights estimated for the farnesoid X receptor (FXR) (0.0%) and peroxisome proliferator-activated receptor gamma (PPARγ) (0.1%) do not allow for the conclusion that the actual biological impact in vivo is correspondingly low. As the Bayesian qAOPE is already described with sufficient mathematical precision by the RAR and PXR signaling pathways, the PPARγ and FXR pathways, due to high collinearity, do not provide any additional predictive power.

We also considered alternative ensemble methods such as Pseudo BMA+ (Bayesian Model Averaging with Bayesian Bootstrap) (Yao et al., 2018) to address the high correlation of the signaling pathways present in the AOP network. Pseudo-BMA+ assumes the existence of a single “true” model, forces a distribution of weights over all pathways and is not suitable for highly collinear models (Yao et al., 2018). In contrast, Bayesian Stacking models the combined prediction distribution by neutralizing the highly correlated, overlapping variance of the individual pathways (Yao et al., 2018), which is inherent in biological networks with converging pathways. Thus, the Bayesian qAOPE approach estimates an optimally parsimonious model suitable for regulatory applications.

The computational effort of the Bayesian qAOPE approach is a limitation. For performing exact Bayesian LOO-CV for each omitted observation, it is necessary to re-estimate large portions of the Bayesian qAOPE using time-consuming MCMC sampling. This computation time is not a major factor for a smaller dataset or for a single active compound, but it may be relevant for high-throughput screening. Thus, it could become necessary to use Pareto Smoothed Importance Sampling (PSIS-LOO-CV) or other approximation methods for larger datasets. The Bayesian qAOPE accurately predicts triglyceride accumulation based on nominal in vitro measurements. Extrapolating these in vitro results to in vivo risk assessment requires the subsequent use of physiologically based toxicokinetic (PBTK) modeling to determine the equivalent oral in vivo concentration, a transformation step already taken for propiconazole (Steinbach et al., 2025).

The latest published Bayesian hierarchical pathways, e.g. the model presented by Zhou and Sahlin (2026), try to integrate pathway dependent effects of the key event relationships and the substance dependent effects into a single model with random effects. The Bayesian qAOPE framework follows a different philosophy: Our framework itself is a chemical agnostic blueprint able to model any complex model including ascending (agonist), descending (antagonistic) and converging relationships thus being applicable to all possible AOP structures. Our estimated pathway ensemble is by design fitted to the toxicodynamics of the tested compound. The Bayesian qAOPE presented here is based on agonist dose-response relationships; future applications of our framework should also include antagonistic relationships like the LXR inhibitor. The application of the Bayesian qAOPE framework to more substances will show if there is the need for an integrated chemical-agnostic model (Zhou & Sahlin, 2026), or if the estimation of a highly accurate chemical-specific pathway ensemble will offer a better way for toxicological predictions in risk assessment.

In conclusion, this study demonstrates that complex, non-linear Bayesian network architectures can identify primary biological drivers with exceptional predictive power. By identifying and consolidating mathematically redundant paths in Bayesian networks through exact LOO-CV Bayesian stacking, the Bayesian qAOPE constitutes a data-driven filter for causal relationships. Our Bayesian ensemble framework offers an easily transferable, modular blueprint for mathematically deciphering complex networks with converging causal paths. This approach is therefore relevant beyond toxicology for disciplines such as epidemiology, systems biology, and pharmacometrics.

